# Unique territorial and compartmental organization of chromosomes in the holocentric silkworm

**DOI:** 10.1101/2023.09.14.557757

**Authors:** J. Gil, E. Navarrete, C. Hockens, N. Chowdhury, S. Abraham, G. Cornilleau, E.P. Lei, J. Mozziconacci, E.J. Banigan, L.F. Rosin, L.A. Mirny, H. Muller, I.A. Drinnenberg

## Abstract

Hallmarks of multicellular eukaryotic genome organization are chromosome territories, compartments, and loop-extrusion-mediated structures, including TADs. However, these are mainly observed in model organisms, and most eukaryotes remain unexplored. Using Hi-C in the silkworm *Bombyx mori* we discover a novel chromatin folding structure, compartment S, which is “secluded” from the rest of the chromosome. This compartment exhibits loop extrusion features and a unique genetic and epigenetic landscape, and it localizes towards the periphery of chromosome territories. While euchromatin and heterochromatin display preferential compartmental contacts, S domains are remarkably devoid of contacts with other regions, including with other S domains. Polymer simulations show that this contact pattern can only be explained by high loop-extrusion activity within compartment S, combined with low extrusion elsewhere through the genome. This unique, targeted extrusion represents a novel phenomenon and underscores how evolutionarily conserved mechanisms—compartmentalization and loop extrusion—can be repurposed to create new 3D genome architectures.

## Introduction

Three-dimensional (3D) genome organization regulates cellular physiology by influencing gene expression ^1–8^, DNA replication ^9,10^, and DNA repair ^11,12^, among other processes ^8,13–16^. This rich, multiscale spatial organization includes chromosome territories (CTs) ^17–19^, chromatin compartments ^20,21^, and extrusion-mediated loops ^22–29^, each of which arises through a distinct physical mechanism ^13,16,30^. These mechanisms simultaneously act on chromatin, raising the possibility that they may interact with each other. Here, we study the genome folding of the silkworm *Bombyx mori* and uncover a novel interplay between two major mechanisms: compartmentalization and loop extrusion.

Chromatin compartmentalization is thought to arise from affinity-mediated phase separation of different types of chromatin. By this mechanism, similar types of chromatin (broadly, euchromatin and heterochromatin) may segregate into spatially distinct compartments ^13,30–33^. In Hi-C and other chromatin contact maps, this effect manifests as a distinctive checkerboarding pattern reflecting preferential interactions between segments of like chromatin state. Studies suggest that compartmentalization maintains epigenetic state, through maintaining its long-term memory ^34,35^ and the co-regulation of epigenetic states between distal loci ^36^.

Loop extrusion is driven by molecular motors belonging to the structural maintenance of chromosome (SMC) complexes, such as cohesin and condensin. These motors bind to chromatin and extrude it into progressively enlarging loops. Cohesin extrudes chromatin and is stalled by barriers, such as CTCF in vertebrates. This process forms TADs in Hi-C maps, which are insulated domains of enriched contacts, as well as dots and stripes ^24,25,37–41^. In vertebrates, the interphase genome is typically fully partitioned into TADs, reflecting a relatively uniform distribution of cohesins across the genome. However, in certain developmental contexts, cohesin can be recruited to specific loci and form a distinct Hi-C contact pattern, termed fountains/jets/plumes ^42–47^. Thus, loop extrusion activity may generate different structures depending on cohesin loading patterns and the presence of extrusion barriers in the surrounding chromatin.

Studies of extrusion in vertebrate systems have often found an antagonistic relationship between loop extrusion and compartmentalization ^26,27,48–50^. Suppressing cohesin activity removes chromatin loops and weakens TADs, while also strengthening compartments ^26,27,48,49^. Increasing cohesin activity by extending its residence time lengthens chromatin loops, strengthens TADs, and weakens compartments ^48–50^. Polymer modeling reveals this antagonism may be caused by cohesin bridging interactions between different types of compartments and increasing chromatin mobility ^32^. However, a recent study demonstrated that loop extrusion is not solely antagonistic to compartmentalization, as stabilized extrusion enhances certain compartments in simulations of mitotic chromosomes ^51^. Thus, the effect of loop extrusion on compartmentalization may depend on the precise activity of loop extrusion in a given context.

While many studies focus on loop extrusion in vertebrates, loop extrusion is well conserved across the kingdoms of life. Indeed, many important insights into vertebrate loop extrusion activity were first established in other organisms. For example, observations of SMC complexes in yeast ^52–55^ and bacteria ^56,57^ were foundational for subsequent studies explaining the nature of SMC-RNA Polymerase interactions in mammals ^58,59^. Similarly, targeted loading of SMC complexes to specific loci in bacteria ^60–63^ preceded the discovery of fountains/jets/plumes in vertebrate cells ^42,43,46^. Therefore, studying diverse organisms beyond vertebrates can reveal new principles of loop extrusion and genome organization.

While a recent study of 3D genomics compared a diverse set of organisms, it excluded species with radically different linear chromosome organization, such as species with holocentric chromosomes ^64^. These have centromeric function distributed along the entire length of each chromosome, unlike most eukaryotes that have a single centromeric region per chromosome. Few studies have investigated the 3D genome organization of holocentric organisms, with the notable exception of *C. elegans* ^65–68^. Thus, studies of such organisms could expand our understanding of the principles of genome architecture.

To this end, we characterize the genome architecture of the holocentric silkworm *B. mori*, which evolved holocentricity convergently with *C. elegans*. By combining Hi-C, epigenetic and transcriptomic profiling, polymer modeling, and Oligopaint FISH, we find a unique genome organization in *B. mori*. This organism exhibits a high degree of chromosome territoriality, three epigenetically distinct compartments, and a nonuniform pattern of loop extrusion activity. Further, we uncover an extreme instance of loop extrusion interfering with compartmentalization. Our analysis and modeling demonstrate that, through targeted loop extrusion, *B. mori* establishes novel, secluded domains that are depleted in chromosome-wide compartment contacts. This unique chromatin structure relies on the interplay of extrusion and compartmentalization and cannot arise by either mechanism alone. Our study thus demonstrates how novel chromatin structures can emerge from new combinations of conserved genome folding mechanisms.

## Results

### Bombyx mori linear genome organization

The *B. mori* genome encompasses 27 autosomes, the Z sex chromosome (Chr01), and the W sex chromosome, which vary in length from approximately 8 to 22 Mb. Genome-wide analysis of GC content revealed a tripartite organization of *B. mori* chromosomes, with a central GC-poor region and two large telomere-proximal GC-rich regions (Figure 1A). This linear organization is reminiscent of that of the nematode *C. elegans* (Figure 1B), another holocentric organism whose chromosomes are divided into centers and arms ^69^. We thus used our GC content track along *B. mori* chromosomes to define arms and center regions for each chromosome (see M&M for detail, Table S1). In *C. elegans*, chromosomal arms are enriched in repetitive DNA, while centers are enriched in genes ^69^ (Figure 1B). In *B. mori*, we found a similar organization with significant, but less distinct, patterns of transposable elements (TEs) and gene coverage (Figure 1B). The similarity in linear genome organization between these two species might reflect convergent evolution between the two holocentric lineages that evolved independently from different monocentric ancestors ^70,71^.

**Figure 1:**
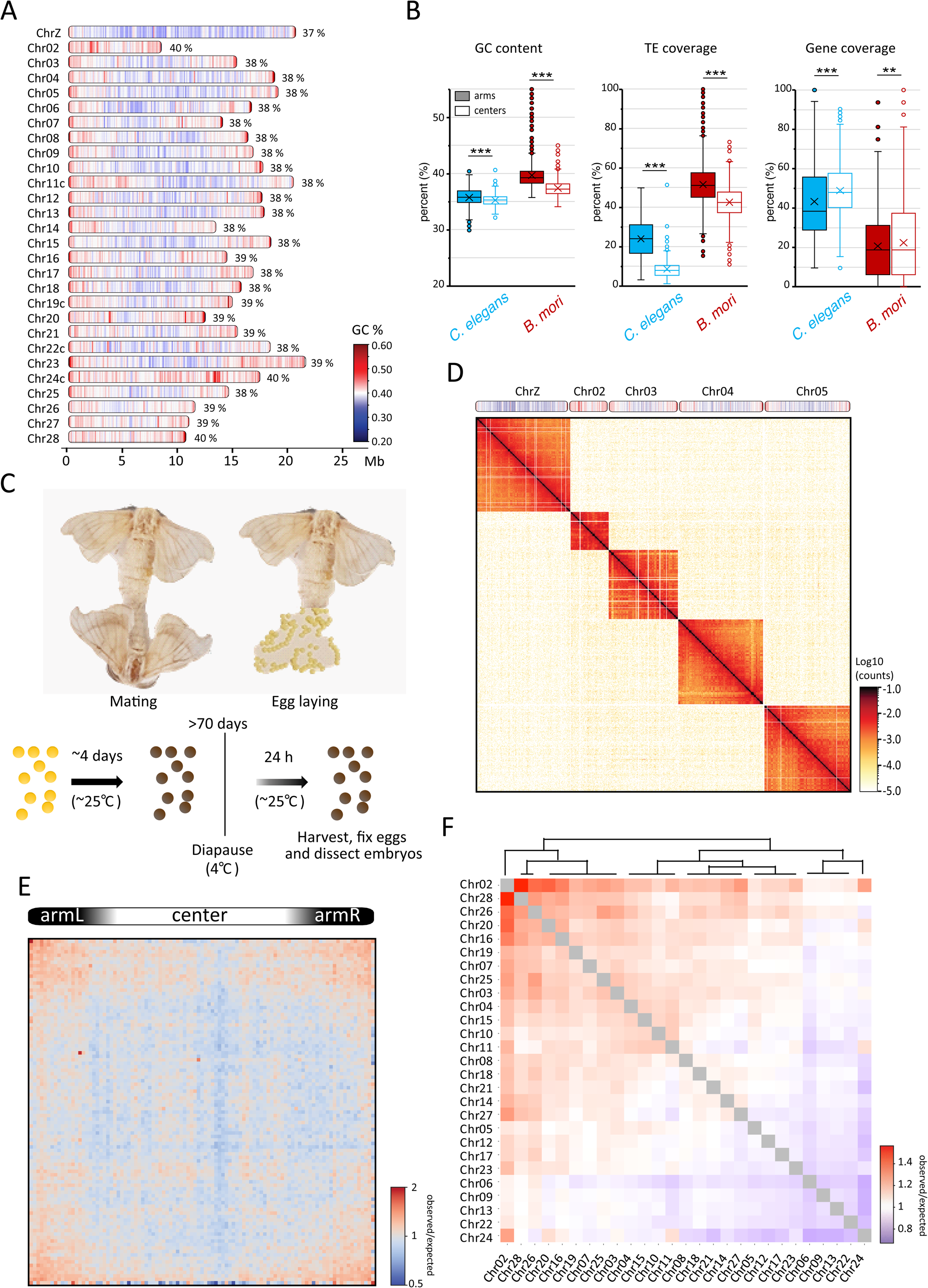
Hi-C of *Bombyx mori* embryos reveals highly distinct chromosome territories. (**A**) Schematics of *B. mori* chromosomes drawn to approximate scale indicated in Mb below. For each chromosome, total GC content in percentage is noted, and local GC content per 100 kb windows is indicated as a color scale from blue (<20%) to red (>60%). A suffix “c” added to a chromosome’s name indicates assembly corrections were made (Figure S9). (**B**) Box plots showing the distribution of GC content, TE coverage, and gene density per 100 kb window in center and arm regions of autosomes in *C. elegans* and *B. mori*. Statistical significance was tested using a Mann-Whitney U-test, **: P-value < 0.05, ***: P-value < 0.005. Box region corresponds to data between the first and third quartile. Lines indicate medians of respective distributions, while crosses correspond to their means. Whiskers extend to the lowest and highest data points, excluding outliers, which are shown as dots. (**C**) Sample generation for Hi-C and ChIP-Seq. (**D**) Contact map at 80 kb bin resolution of five selected *B. mori* chromosomes (*on top*). (**E**) Average inter-chromosomal (*trans*) ^observed^/_expected_ matrix for all scaled *B. mori* chromosomes, computed at 40 kb bin resolution. (**F**) Heatmap illustrating ^observed^/_expected_ inter-chromosomal contact frequencies as a divergent color scale from blue to red. Chromosomes have been clustered and ordered to reflect similar contact patterns.

### Chromosomes in B. mori embryos form highly segregated territories

We generated Hi-C datasets from whole *B. mori* embryos at three different embryonic post diapause time points (2, 24, and 48 hours after diapause release) and one adult stage from the p50 reference strain. For most of our analysis, we focused on the 24-hour post-diapause timepoint (PD-D2) (Figure 1C), for which we confirmed that a proportion of cells have re-entered the cell cycle and are thus no longer arrested in G2 (Figure S1) ^72^.

The PD-D2 Hi-C contact pattern across all 28 chromosomes revealed very sparse inter-chromosomal contacts (Figure 1D). This is consistent with recent DNA FISH data that revealed that *B. mori* chromosomes are tightly folded and occupy distinct CTs ^73^. Despite the similarity in linear genome organization between *B. mori* and *C. elegans*, clustering between centers and arms of different chromosomes, easily seen on *C. elegans* contact maps ^65,66^, is not evident in *B. mori*. Nevertheless, drawing the average *trans* contact matrix reveals enrichments between large sub-telomeric regions, indicating some degree of telomere or arm clustering between chromosomes in *B. mori* (Figure 1E).

Despite the sparsity of inter-chromosomal contacts, we tested whether known patterns of sub-nuclear positioning of chromosomes seen in other organisms are present in *B. mori*. In humans, small, gene-rich chromosomes have been shown to preferentially interact with each other and localize more centrally within the nucleus ^20,74–76^. A length-dependent contact preference can also be identified among *B. mori* chromosomes, with a group of small chromosomes (Chr02, 28, 26, 20, and 16) having the highest average inter-chromosomal contact frequency (Figures 1F, S2A). However, the correlation between inter-chromosomal contacts and gene density is mild (Figure S2B). This might be due to the lower variation in length and gene content among *B. mori* chromosomes compared to human chromosomes. We also noticed a positive correlation between inter-chromosomal contacts and GC content (Figure S2C), but whether GC content contributes to observed contact preferences is unclear.

Our data reveal conserved principles in chromosome organization in *B. mori*, such as the formation of CTs and mild length-dependent inter-chromosomal contact preferences. In contrast to other organisms, however, inter-chromosomal contacts are not correlated with gene density, and *B. mori* chromosomes make very limited inter-chromosomal contacts overall, highlighting the remarkably strong CTs in this organism.

### B. mori chromosomes are organized in three chromatin compartments

We next investigated compartmentalization of *B. mori* chromosomes. Many regions of the Hi-C maps displayed a characteristic checkerboard pattern of alternating regions of two types, displaying enriched homotypic (self-to-self) and depleted heterotypic (self-to-other) contacts (Figure 2A, left and middle panels), seen in all other animals ^8,30,77–81^. However, unlike Hi-C maps from any other organism, we unexpectedly found large chromatin segments that do not display checkerboard patterning. Instead, these regions engage in a high degree of short-range contacts but few long-range contacts with any other chromosomal segments (Figure 2A, left and right panels). Therefore, these regions are atypical for compartments, but also appear to be different from TADs, which are described in vertebrates and exist within checkerboarding compartments.

**Figure 2:**
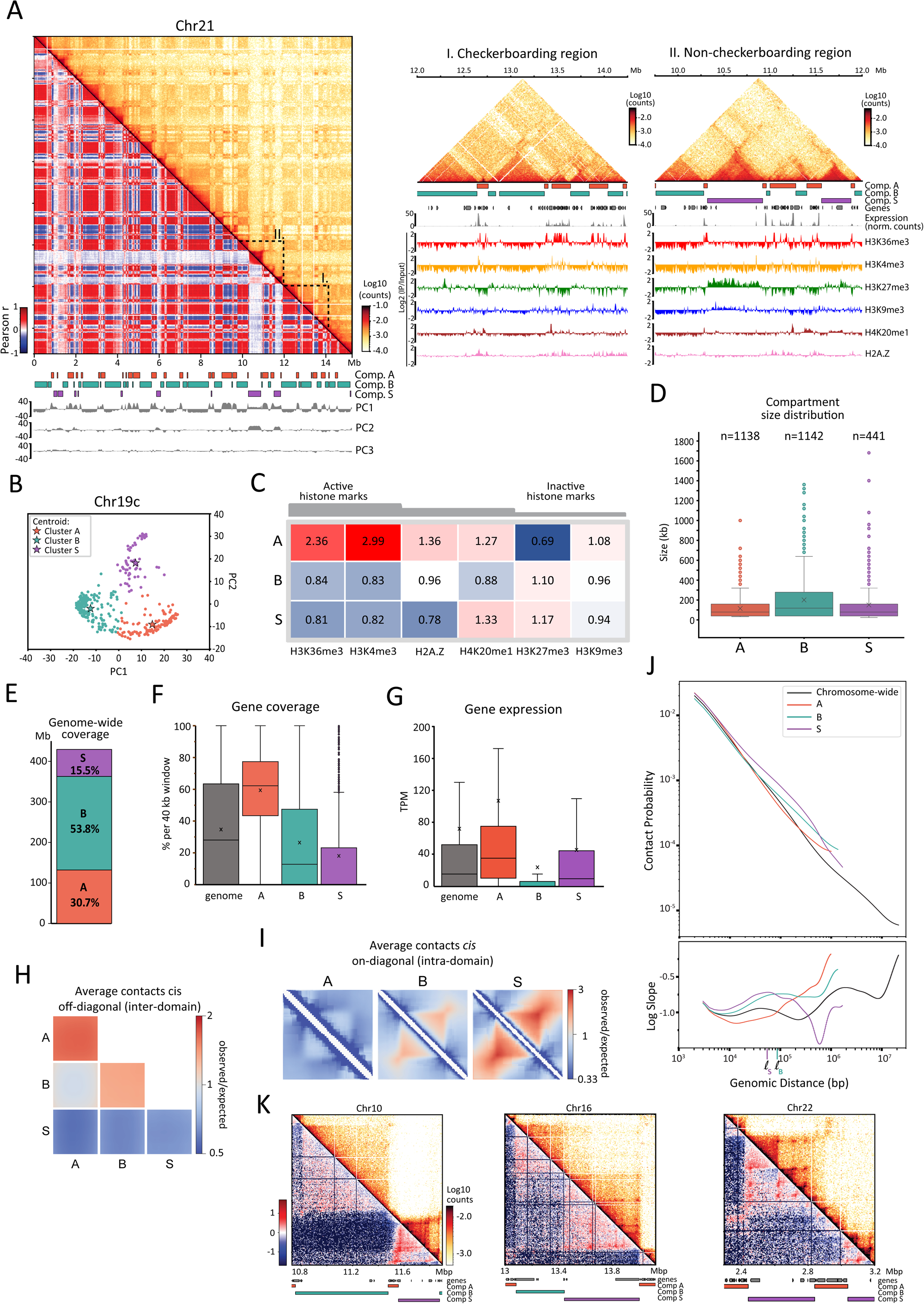
Hi-C compartment analysis reveals a novel and secluded chromatin folding structure. (**A**) *Left*: top diagonal represents the iterative-corrected Hi-C contact map of Chr21 at 40 kb bin resolution, and bottom diagonal is the corresponding Pearson correlation matrix. Below the matrix are the gene track (black), color boxes indicating the locations of domains of compartment A (orange), B (turquoise), and S (purple) and tracks of the first three principal components (gray) computed from the correlation matrix. *Right*: close-up on regions highlighted on the left, I: checkerboarding region, II: non-checkerboarding region. Below matrices are tracks for compartments and genes as in the left as well as corresponding RNA-seq at 1 kb resolution and ChIP-Seq tracks for various histone marks at 25 bp resolution. (**B**) Example of clustering along PC1 and PC2 of Pearson correlation matrix for Chr19c. Clusters called using k-means function of scikit-learn are shown in different colors. For each cluster, the centroid and the corresponding assignment to A, B, and S compartments is indicated. (**C**) Heatmap showing genome-wide enrichment of assayed histone marks across compartment A, B, and S. Enrichment values correspond to median of IP/input ratios at 40 kb resolution normalized to the genome-wide median for each mark. (**D**) Box plots showing the size distribution of A, B, and S domains. Boxed regions correspond to data between the first and third quartile. Lines indicate the medians of respective distributions, while crosses correspond to their means. Whiskers extend to the lowest and highest data points, excluding outliers, shown by dots. (**E**) Bar graph showing relative genomic coverage of A, B, and S. (**F**) Box plots showing distribution of gene coverage per 40 kb window of the whole genome (n = 11,143) or within A (n = 3,294), B (n = 5,772), and S (n = 1,670). Line indicates median of distribution while cross corresponds to mean. Whiskers extend to lowest and highest data points, excluding outliers, which have been removed. All distributions in each compartment for all features are significantly different from each other and from the genome-wide distribution by the Mann-Whitney (95%, two-tailed) U-test. (**G**) Box plots showing range of gene expression (in TPM) (whole genome: n = 13,869, in A: n = 7,404, in B: n = 4,500, in S: n = 608). (**H**) Average *cis* off-diagonal (inter-domain) ^observed^/_expected_ contact enrichment plots (log scale) within and between all rescaled A, B, and S compartments. (**I**) Rescaled average *cis* on-diagonal (intra-domain) ^observed^/_expected_ contact enrichments (log scale) for the three compartment types. (**J**) *Top:* Contact probability as a function of genomic distance, P(*s*), curve for the PD-D2 Hi-C data (*s*). *Bottom*: Log-derivative of contact probability as a function of genomic distance. Curves represent averages of either chromosome-wide (gray) or for contiguous segments of a given compartment type (colored). Estimates of the average loop size in S (ℓ_S_) and in B (ℓ_B_), corresponding to the peaks in the log-derivative curves, are labeled on the x-axis. (**K**) Regions of the PD-D2 Hi-C maps (top diagonal) along with the corresponding Pearson correlation matrix (bottom diagonal) displaying characteristic features of loop extrusion at 5 kb resolution, with compartment and gene tracks below.

To identify which chromosomal loci engage in this novel interaction pattern, we applied an approach similar to a recently developed sub-compartment detection algorithm ^82^. From the *cis* Hi-C maps, we k-means clustered the leading principal components of the Pearson-correlated matrices, revealing groups of loci with similar contact profiles (Figures 2A, and Table S2). This strategy enabled us to define three main clusters along each chromosome, which we then unified across the genome based on epigenetic composition (Materials and Methods).

We profiled active (H3K36me3, H3K4me3) and silent (H3K27me3, H3K9me3) histone marks by ChIP-seq in the PD-D2 embryonic stage of *B. mori*. We also included H2A.Z, a histone variant associated with transcriptional control ^83^, which is enriched in a sub-compartment that shows an attenuated checkerboarding pattern in the cancer cell line HCT116 ^82^. Additionally, we included H4K20me1, a mark associated with centromeric nucleosomes in vertebrates ^84^, as well as a variety of processes including transcriptional regulation, chromosome replication and segregation, DNA damage response, and chromosome compaction ^85,86^.

Consistent with the compartmentalization of euchromatin and heterochromatin observed in other animals, we found that the two checkerboarding clusters were enriched in active (“compartment A”) and inactive (“compartment B”) histone marks (Figures 2A, 2C, S3A, and S3B). The third cluster captured the novel, non-checkerboarding regions. We termed this cluster “compartment S,” reflecting its secluded, spatially segregated behavior observed in the Hi-C maps. Compartment S displayed an epigenetic signature consistent with silenced chromatin, though it uniquely differed from compartment B due to its enrichment in H4K20me1.

Domains, which we define as contiguous segments of the same compartment type, of A, B, and S are heterogeneous in length and cover different fractions of the genome (Figures 2D, 2E). Overall, A and S domains are smaller, with a median size of 80 kb, while B domains are generally larger, with a median size of 120 kb. A and B compartments cover approximately 50 and 30% of the genome, respectively, while S covers 15%, with 5% of the genome not assigned to any of the three clusters. Consistent with our description of *B. mori* linear genome organization, we found domains of compartment A are also enriched along chromosome centers, while those of compartment B are enriched along the arms (Figure S3D). Domains of compartment S are distributed throughout the genome, lacking preferential positioning at either chromosome centers or arms (Figure S3D and S3C for the distribution of S domains).

We next investigated whether compartment S is distinct from compartments A and B with respect to genetic features. We found that compartment S, like compartment A, has a lower GC and transposable element content compared to the whole genome (Figure S3E). Despite these similarities to A, compartment S corresponds to gene-poor regions and is even more depleted in genes than compartment B (Figure 2F). However, unlike B, S does not appear to be a repressive compartment; the expression levels of the few genes that are located within S are in the range of the genome-wide distribution. Out of the 608 genes within S, about half (300) are expressed at the PD-D2 stage (TPM > 10) (Figure 2G and Table S3). Gene ontology analyses revealed genes within S are enriched in DNA-binding and transcription regulation processes (Table 1). Given that one of the largest S domains in the genome corresponds to the Hox cluster, which is expressed in the PD-D2 stage, we tested for an enrichment of homeotic and Polycomb group (PcG) response genes (Figure S4). Using previously published datasets ^87,88^, we generated a list of 399 homeotic and PcG response genes in *B. mori* and assigned them to A, B, or S (Table S4, Materials and Methods). We did not find an enrichment of homeotic or PcG response genes in compartment S but rather in A (Table S4).

**Table 1:** Molecular Function enrichment of *B.mori* genes in S.

| Pathway | GO ID | Enrichment FDR | Number of genes in category | Total number of pathway genes | Fold Enrichment |
|---|---|---|---|---|---|
| DNA binding | GO:0003677 | 6.10E-23 | 66 | 592 | 4.45 |
| Nucleic acid binding | GO:0003676 | 7.43E-14 | 89 | 1461 | 2.43 |
| DNA-binding transcription factor activity | GO:0003700 | 3.18E-13 | 31 | 220 | 5.63 |
| Transcription regulator activity | GO:0140110 | 5.13E-12 | 32 | 262 | 4.88 |
| Sequence-specific DNA binding | GO:0043565 | 7.10E-08 | 19 | 139 | 5.46 |
| DNA-binding transcription factor activity, RNA polymerase II-specific | GO:0000981 | 2.811E-06 | 13 | 80 | 6.49 |
| Nuclear receptor activity | GO:0004879 | 4.05E-03 | 4 | 12 | 13.32 |
| Ligand-activated transcription factor activity | GO:0098531 | 4.05E-03 | 4 | 12 | 13.32 |
| Steroid hormone receptor activity | GO:0003707 | 1.22E-02 | 4 | 16 | 9.99 |
| Signaling receptor binding | GO:0005102 | 2.77E-02 | 7 | 67 | 4.17 |

We conclude that *B. mori* chromosomes segregate into three chromatin compartments. In addition to the well-conserved compartments A and B*, B. mori* chromosomes contain a third compartment type, compartment S, which has an unprecedented pattern of contact. S exhibits a unique epigenetic landscape and covers about one-sixth of the *B. mori* genome at the PD-D2 time point. While S largely encompasses gene-poor regions, many genes located within S are transcriptionally active.

### Compartment S exhibits novel contact patterning and key evidence of loop extrusion activity

To formalize our initially observed contact patterning for compartments A, B, and S, we calculated the average compartment contact strengths for all domains on the same chromosome. For A and B, we found enriched homotypic (A-A and B-B) and depleted heterotypic (A-B) interactions, confirming that A and B are compartmentalized (Figure 2H). Further, we found that domains of S were depleted in contacts with all other compartments. These formed smooth lanes of contact depletion, such that A-S, B-S, and S-S (other S domains) were all depleted at similar strengths (Figure 2H). In stark contrast to this finding, we found contacts for pairs of loci within the same, contiguous S domain (on-diagonal) were strongly enriched (Figure 2I). To our knowledge, this unexpected discordance between on- and off-diagonal Hi-C contacts has not been observed in other organisms, and it raises the question of whether this pattern arises from a previously undescribed mechanism.

In vertebrates, cohesin extrudes loops resulting in domains of local contact enrichment known as TADs and decreased compartmentalization ^27,32,49,50^. Given both the high contact enrichment within S domains (Figure 2I) and their lack of checkerboard patterning (Figure 2A and H), we hypothesized that S domains might represent regions of high loop extrusion activity. We searched for evidence of loop extrusion activity in the PD-D2 Hi-C data. Key established indicators of this activity are (1) the shape of the contact frequency *P(s)* as a function of genomic separation *s* ^19,27,48,89,90^ and (2) the presence of specific Hi-C patterns, such as insulated domains, dots, and stripes, which also require extrusion barriers ^24,26,27,41,91,92^.

In the chromosome-wide *P(s)* curve (Figure 2J), we observe a characteristic “shoulder” of increased contacts and a corresponding peak in the log-derivative of *P(s)*, which are indicative of loop extrusion ^19,27,48,89,93^. The peak in the log derivative plot represents the average loop size ^48,90^, which in this case is approximately 40-60 kb. We next computed these contact frequency curves separately for continuous segments of each compartment type (Figure 2J). Consistent with our hypothesis, the *P(s)* shoulder is most prominent for compartment S and indicates an average loop size of 40-60 kb. The *P(s)* curve in compartment B also suggests some loop extrusion activity, though extrusion in B is likely sparser with larger loops, as suggested by the smaller height and right-shift of the peak. Analysis of *P(s)* therefore indicates that loop extrusion is active in *B. mori* and is enriched in S domains.

In further support of loop extrusion in *B. mori*, we found many genomic regions with features of barrier-restricted extrusion (dots and stripes). By visual inspection, we often found such features nested within each other inside and at the edges of S domains (Figure 2K).

The Hi-C data strongly suggests that loop extrusion occurs on *B. mori* interphase chromosomes and that it is much more prominent in S domains. This preferential enrichment of extrusion differs from previous observations in vertebrates, which raises the mechanistic question of whether it is responsible for the nontrivial patterns of S contacts in *B. mori*.

### Targeted loading of loop extruders to S domains can reproduce S compartment Hi-C features

Given the novel and stark contact pattern of compartment S (Figure 3A), we employed polymer modeling to systematically study which mechanisms underlie its formation. We sought to reproduce the following three Hi-C features that together make compartment S unique (Figure 3A): (*i*) contacts are enriched within contiguous, on-diagonal S domains (locally enriched, S_intra_); (*ii*) each S domain is depleted in off-diagonal, compartment contacts with other S domains (distally depleted, S-S_inter_); (*iii*) the depletion of contacts between S domains and the rest of the chromosome is homogenous (smooth lanes of contact depletion, S-S_inter_ ≈ S-A ≈ S-B). These features are the hallmarks of compartment S and have not been previously observed together or at all.

**Figure 3:**
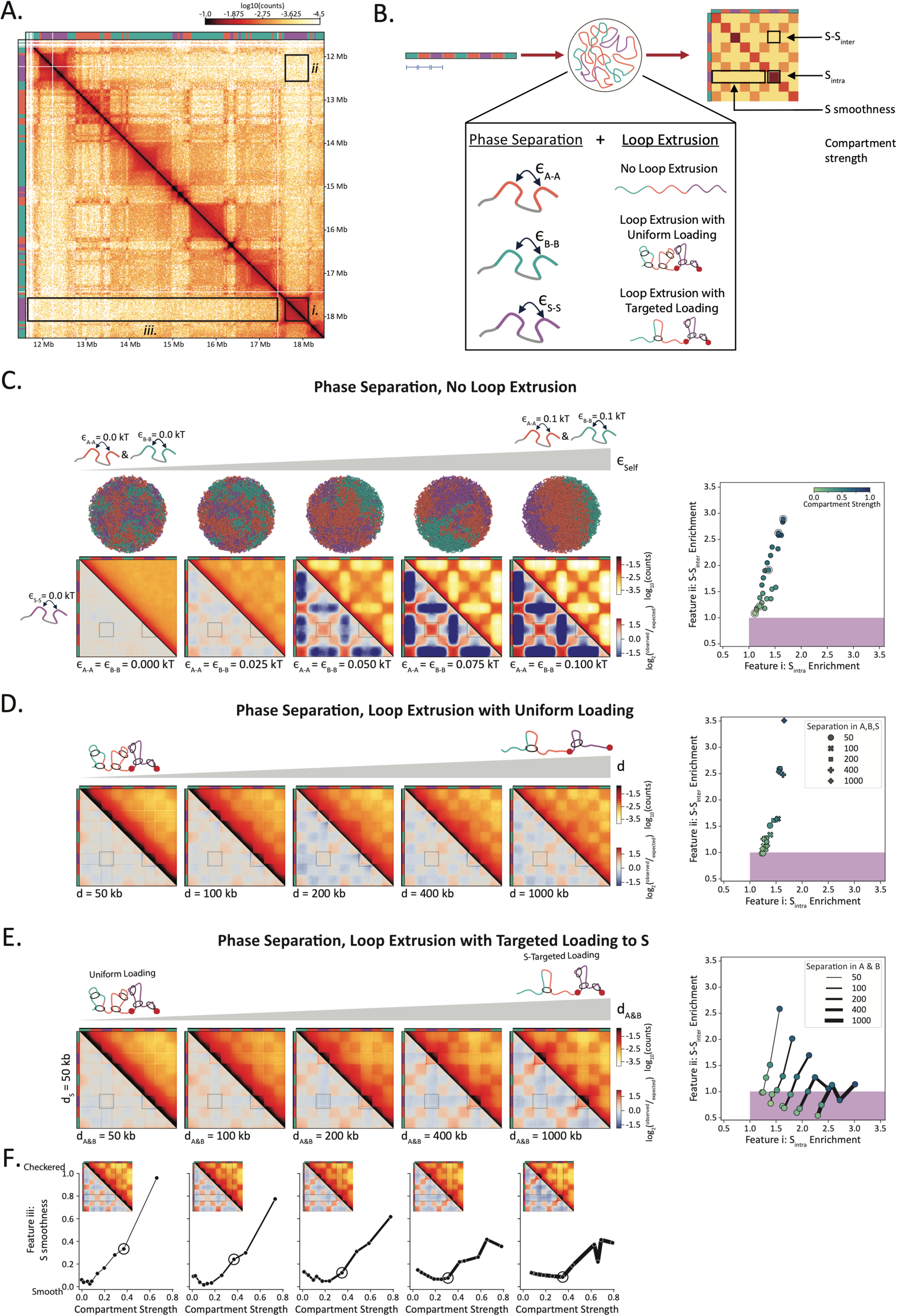
Mechanistic polymer modeling reveals S chromatin organization can arise from targeted loop extrusion in S chromatin but not from other known genome folding mechanisms. (**A**) Hi-C map and sub-compartment annotations highlighting the three distinct features of S compartments: *i*) on-diagonal contact enrichment within S domains, *ii*) off-diagonal contact depletion between different S domains, and *iii*) smooth depletion of contacts between S and the rest of the chromosome. Hi-C data corresponds to chromosome 22: 11,625,000-18,435,000 from PD-D2 samples mapped with distiller (see Methods). (**B**) Illustration of the approach to testing different mechanistic models of S compartment formation. Polymer segments are assigned to compartments A, B, or S, which specifies their pairwise affinities. One model without loop extrusion and two models with loop extrusion are considered. Simulations are evaluated based on whether their contact maps display features *i*, *ii*, and *iii*. Compartment strength is also measured. (**C**) Three-species phase separation modeling results. *Left*: sampled conformations and corresponding Hi-C maps of chromosome models with increasing phase separation driven by increasing both A-A and B-B affinities, ɛ_A-A_ and ɛ_B-B_ (from left to right). *Right*: scatter plot of ^observed^/_expected_ contact enrichments of S features *i* and *ii* for all simulated A-to-A and B-to-B affinities. Purple boxed region reflects the broadest value range for S-like features. Data points corresponding to chromosome models shown on the left are circled in the scatter plot. (**D**) Phase separation and uniform loop extrusion with barriers modeling results. *Left:* Hi-C maps of models with increasing chromosome-wide separation, d, of loop extruders. Phase separation was induced in all maps on the left by setting ɛ_A-A_ = ɛ_B-B_ = 0.05 kT. *Right:* scatter plot of ^observed^/_expected_ contact enrichments of S features *i* and *ii* for models varying in extruder separations, A-A, and B-B affinities, with points colored by compartment strength as in (**C**). For all models shown, processivity was fixed to 100 kb and S-to-S affinity, ɛ_S-S_, set to 0.0 kT. (**E**) Phase separation and loop extrusion with targeted loading to S modeling results. *Left*: Hi-C maps of models with increasing separation of loop extruders in A and B (d_A&B_), while extruder separation in S (d_S_) was fixed at 50 kb. Phase separation was induced in all maps on the left by setting ɛ_A-A_ = ɛ_B-B_ = 0.05 kT. *Right*: scatter plot of ^observed^/_expected_ contact enrichments of S features *i* and *ii* for a range of A-to-A and B-to-B affinities and separations in A and B (d_A&B_), with d_S_ fixed to 50 kb. Data points sharing d_A&B_ (i.e. varying ɛ_A-A_ and ɛ_B-B_) were connected by line, with the value of d_A&B_ reflected in line thickness. The line corresponding to d_A&B_ = 50 kb reflects the uniform loading scenario. (**F**) Impact of S-targeted extruder loading on the smoothness of S compartment contact depletion (feature *iii*). Smoothness was computed as the standard deviation of mean observed/expected S-A, S-B, and S-S_inter_ contacts, with lower y values corresponding to smoother compartment patterning. Data points with compartment strengths of approximately 0.4 are circled and corresponding Hi-C maps are shown in the insets. Separation in S (d_S_) was fixed at 50 kb.

In addition to its unique loop extrusion activity, we found S exhibits a unique epigenetic composition, suggesting it may phase separate from compartments A and B. Because loop extrusion and phase separation independently drive chromatin folding, we asked which of these processes, either individually or in combination, are responsible for generating S features. We therefore tested three major classes of mechanistic models (Figure 3B). We first tested models in which chromatin organization was solely driven by phase separation. We then tested models with uniform loop extrusion and extrusion barriers at the edges of S domains, akin to loop extrusion behavior that generates TADs in vertebrates. These first two classes represent mechanisms of genome folding described in other animals. Finally, we tested models in which loop-extruding complexes preferentially accumulate in S over A and B.

We performed polymer simulations for each class of model, using simplified (chromatin) polymer chains with eight equally sized segments of A, B, and S monomers. Monomer identity was used to define compartment-specific affinity (**ε**) and loop extrusion properties (Figure 3B). For each class of model, we tested a wide range of A/B compartment strengths to recapitulate the A/B compartments observed in the experimental data.

#### Phase separation without loop extrusion

We investigated whether affinity-mediated phase separation, which drives the formation of chromatin compartments, could recapitulate the features of S. Experimental Hi-C maps revealed clear compartmentalization of A and B regions (i.e. A/B checkerboarding), whereas S lacked such compartment patterning. This led us to first ask whether S could emerge if A and B chromatin, but not S, exhibited self-affinity.

To test this in our polymer simulations, we systematically varied per-monomer homotypic affinities of A-to-A and B-to-B monomers to obtain a broad range of compartment strengths (Figure 3C). As expected, applying A-to-A and B-to-B affinities produced the desired checkerboard patterning of A and B compartments, consistent with their phase separation. However, S also checkerboarded, so none of the simulated models were able to replicate the distal features of S (features *ii* and *iii*). While we observed a subset of models with on-diagonal enrichment of S_intra_ contacts (feature *i*), it was always accompanied by off-diagonal enrichment of S-S_inter_ contacts—contrary to the depletion required for feature *ii*.

To further understand this relationship, we quantified contact enrichment and depletion from ^observed^/_expected_ (Materials and Methods) Hi-C maps for each model. We assessed whether the Hi-C maps were consistent with the first two S features (S_intra_ and S-S_inter_) using the most lenient criteria: enrichment was defined as an average ^observed^/_expected_ pixel strength greater than one, while depletion was defined as ^observed^/_expected_ less than one. This analysis confirmed that none of the phase separation models accurately captured the experimentally observed features of S. Enrichment of S_intra_ (feature *i*) and depletion of S-S_inter_ (feature *ii*) did not co-occur because contact enrichment within contiguous domains (S_intra_, feature *i*) was correlated with contact enrichment between S domains (S-S_inter_, opposite of feature *ii*). By quantifying compartment strength from each Hi-C map (color scale in Figure 3C, right), we found that both S_intra_ and S-S_inter_ enrichment increased as chromatin compartmentalization strengthened. As evident in the simulated Hi-C maps, models with sufficiently high A-to-A and B-to-B affinities to produce A/B checkerboarding always resulted in S-S_inter_ enrichment and S checkerboarding (Figure 3C, Figure S5A). Together with polymer conformations and the Hi-C maps (Figure 3C, left), these results indicate that phase separation of A and B chromatin induces phase separation of S chromatin, even in the absence of S self-attraction.

To confirm that phase separation of S leads to these undesired, conventional compartment patterns in S (enriched S-S_inter_ and S checkerboarding), we introduced self-attraction to S monomers and repeated the A-to-A and B-to-B affinity sweep. As expected, S-to-S self-attraction resulted in increased S_intra_, S-S_inter_, and S checkerboarding (Figure S5B). We therefore conclude that phase separation does not drive S-like organization.

#### Uniform loop extrusion with A and B phase separation

Informed by evidence of loop extrusion activity in the PD-D2 Hi-C maps (Figure 2J and 2K), we tested whether loop extrusion plays a role in S compartment formation. Given that TAD formation via the loop extrusion process results in domain enrichment and compartment suppression ^27,32,49,50^, we hypothesized that loop extrusion might promote S_intra_ contacts (feature *i*, domain enrichment) while suppressing both S-S_inter_ contacts and the checkering of S (features *ii* and *iii*, compartment suppression). We therefore tested whether S domains are akin to TADs in vertebrates, formed by uniformly loaded, genome-wide loop extruders that are stalled by barriers.

In these models, loop extruders randomly bind to the chromatin fiber and progressively extrude the chromatin polymer into loops. Extrusion proceeds until the extruder unbinds or stalls. Given the evidence for extrusion barriers in and at the edges of S domains (Figure 2K), we placed loop-extrusion-stalling sites at the ends of S domains. We then varied the relevant simulation parameters: A and B homotypic affinities, mean separations between extruders (d), and extruder processivity (λ, the average loop size extruded by an unobstructed extruder). We assumed S monomers lacked preferential attractions to other S monomers.

As evident in the resulting contact maps, these models cannot generate S compartments (Figure 3D, left). We observed on-diagonal extrusion domains in S, with barrier features visually similar to those in the experimental Hi-C data (feature *i*). However, we did not observe off-diagonal depletion of contacts (feature *ii*). Like the phase-separation-only models, this class of models failed to achieve the desired compartment-level organization. Though loop extrusion counteracted compartmentalization, this antagonistic effect was not restricted to S; when compartment features (checkering off-diagonal contacts) were suppressed in S, they were likewise suppressed in A and B. Conversely, when compartment checkerboarding was present in A and B, it was also present in S.

Similar to the phase-separation-only models, upon quantifying Hi-C contact enrichments, we found that S_intra_ and S-S_inter_ contacts are highly correlated and are largely reflective of compartment strength (Figure 3D, right). Consistent with the antagonism between loop extrusion and compartmentalization, stronger extrusion (smaller separations and/or larger processivities) demands higher affinities to achieve the same level of compartmentalization (Figure S5C). While two models narrowly satisfied the quantitative criteria for features *i* and *ii*, we find that their Hi-C maps visually lacked compartmentalization of any type (S-S_inter_ ≈ A-A ≈ B-B contacts, all low), as expected for the high density of extruders (d = 50 kb) (Figure 3D, left). We conclude that the uniform loading model is unable to reliably generate S compartments. Therefore, the presence of S features strongly indicates that some aspect of the loop extrusion process in *B. mori* differs from that of vertebrates.

#### Loop extrusion with targeted loading in S and A/B phase separation

To preferentially suppress compartmentalization of S (off diagonal S features - *ii* and *iii*) while preserving them in A and B, we hypothesized that loop extrusion may be targeted to S. We therefore tested a new class of models in which loop extruders were more likely to load in S than in A and B, resulting in lower mean separation of extruders in S (d_S_) than in A and B (d_A&B_). Our hypothesis of more extruders in S would lead to smaller loops in S, consistent with our observations from the Hi-C data (Figure 2J).

Models with targeted loading to S reliably produced S-like features in their Hi-C maps, unlike the previous two classes of models. Weak extrusion in A and B allowed A/B compartments to form, while strong extrusion in S both compacted S domains (S_intra_ enriched, feature *i*) and inhibited S-S_inter_ compartment contacts (feature *ii*) (Figure 3E, left). Additionally, Hi-C maps with weaker extrusion in A and B had smoother lanes of S depletion (feature *iii*).

Analyzing the Hi-C maps quantitatively showed that targeting loading of extruders to S alters the balance between S_intra_ and S-S_inter_ contacts (Figure 3E, right). As extrusion in A and B weakened (d_A&B_ increased, increased relative targeting), both features *i* and *ii* began to look more S-like: S_intra_ enrichment strengthened (gain of feature *i*) and distal S-S_inter_ enrichment was lost (gain of feature *ii*). Therefore, unlike the phase-separation-only and uniform loop extrusion models, targeting extrusion to S domains both compacts S and secludes them from contacting each other.

Additionally, models with stronger targeted loading produced S features more robustly. First, decreasing extrusion in A and B (increasing d_A&B_) allowed features *i* and *ii* to co-exist across a broader range of compartment strengths (Figure 3F). Furthermore, S features persist upon the introduction of S-to-S affinity comparable to those of A-to-A and B-to-B (Figure S5D).

We next measured feature *iii* as the standard deviation of average ^observed^/_expected_ S-A, S-B, and S-S_inter_ contact frequencies. Smoother lanes of depletion (off-diagonal S contacts) result in smaller standard deviations. In the uniform loading model, once compartment features began to emerge (compartment strength > 0.1), the standard deviation increased (feature *iii* disappeared), indicating a less smooth pattern of inter-compartmental contacts. However, as extrusion in A and B decreased (more targeted loading, i.e. d_A&B_ increased), higher compartment scores could be tolerated before S smoothness was lost. We found that the gain in smoothness (small standard deviation) due to targeted loading is largely driven by the suppression of S-S_inter_ (feature *ii*) as well as additional contributions from A/B compartmentalization (Figure S5E).

Thus, preferential loading of loop extruders to S in combination with self-affinities of A and B reliably produces the patterns of *B. mori* Hi-C maps. The latter compartmentalize the genome, while loop extrusion targeted to S generates the unique features of S compartments.

### The different features of S rely on distinct aspects of targeted extrusion and compartmentalization

To understand how these principles generate the specific folding patterns of *B. mori* chromosomes, we developed a quantitative chromosome-scale model employing the principles from the minimal model study. Focusing on a 6.5 Mb segment of chromosome 15 (Figure 4A), we assigned compartment identities to the simulated chromosome based on those from the PD-D2 Hi-C maps. To compare models to experimental Hi-C, we computed ^observed^/_expected_ contact enrichments between all types of compartments, partitioning S-S contacts into S-S_inter_ and S_intra_. Models varied in five parameters: A-to-A affinity, B-to-B affinity, extruder processivity (λ), the average separation between extruders inside S (d_S_), and the average separation between extruders outside of S (inside A and B regions, d_A&B_).

**Figure 4:**
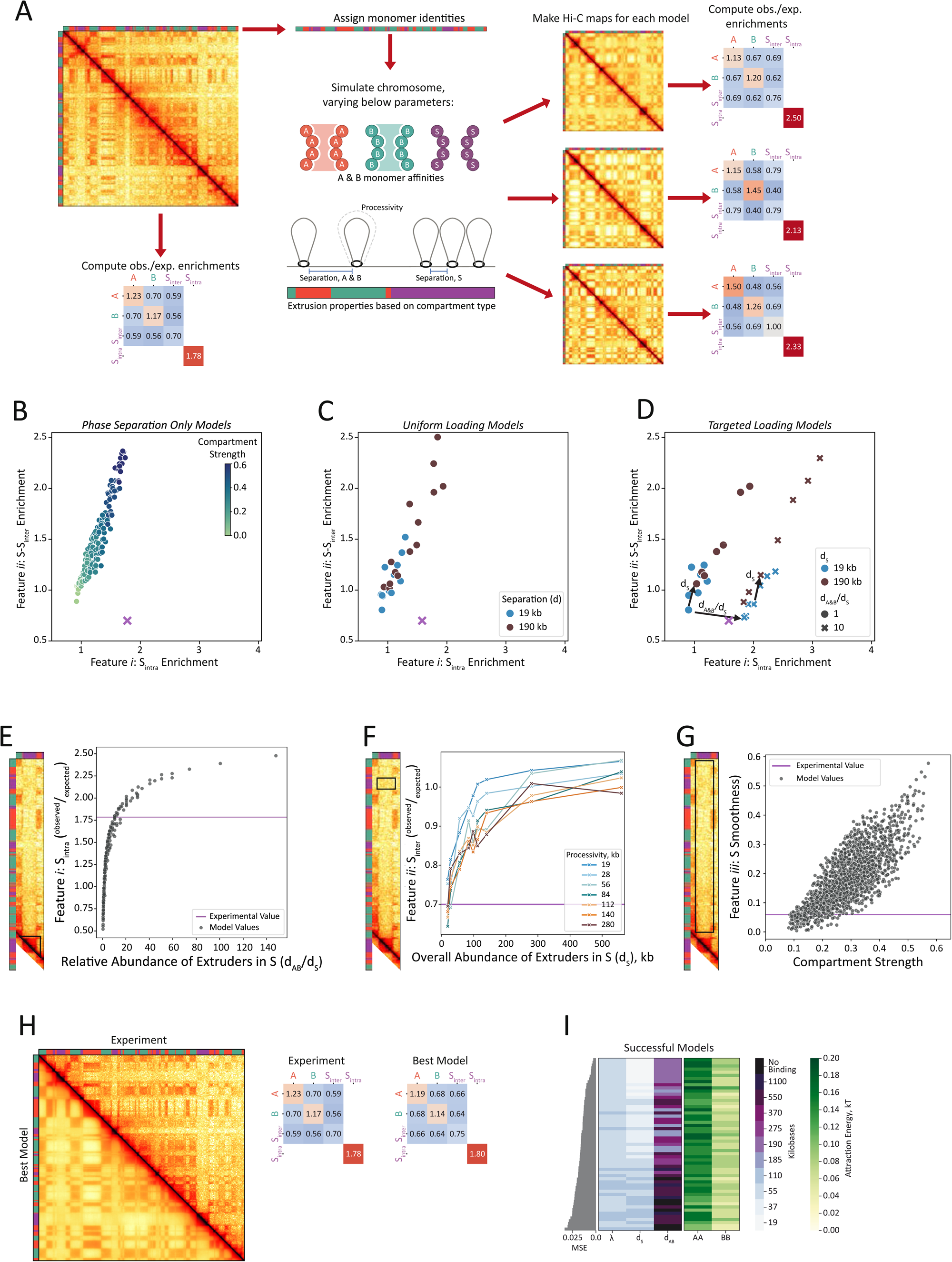
Chromosome-scale polymer modeling reveals how different features of S arise. (**A**) Approach to generating chromosomal polymer models of *B. mori*. Locus corresponds to chromosome 15: 0-6,500,000 mapped with distiller (see Methods). Polymer segments are assigned to compartments A, B, or S based on sub-compartment assignments of the region. Simulations are performed for three different phase separation and/or extrusion models. ^Observed^/_expected_ contact enrichments are computed for different intra- and inter-compartment interactions for comparison to the experimental Hi-C data. (**B**) Scatter plot of ^observed^/_expected_ contact enrichments of S features *i* and *ii* from chromosome-scale polymer modeling of three-species phase separation and no loop extrusion over a range of different A-to-A and B-to-B affinities. The corresponding S_intra_ and S-S_inter_ values from chromosome 15: 0-6,500,000 of the PD-D2 Hi-C data are shown as a purple x. (**C**) Scatter plot of ^observed^/_expected_ contact enrichments of S features *i* and *ii* from chromosome-scale polymer modeling of three-species phase separation and loop extrusion with uniform loading for two different separations, d = 19 or 190 kb, and two processivities, λ = 19 and 190 kb, over a range of A-to-A and B-to-B affinities and extrusion processivities. (**D**) Scatter plot of ^observed^/_expected_ contact enrichments of S features *i* and *ii* from chromosome-scale polymer modeling of three-species phase separation and loop extrusion with targeted loading to S for two different d_S_ and d_S_/d_A&B_ ratios over a range of different interaction affinities. (**E**) ^Observed^/_expected_ contact enrichments of S_intra_ plotted as a function of relative abundance of loop extruders (d_A&B_/d_S_) for a series of chromosome models with λ, d_S_, and d_ACB_ varied. The experimental value of S_intra_ is shown as a purple line. A-to-A and B-to-B affinities were fixed at 0.12 and 0.04 kT, respectively. (**F**) ^Observed^/_expected_ values of S-S_inter_ plotted as a function of separation between loop extruders in S (d_S_) for a series of chromosome models. Each curve represents a series of models, which share loop extruder processivity. All models lacked extrusion in A and B chromatin and shared homotypic attractions of ɛ_A-A_ = 0.12, ɛ_B-B_ = 0.04, and ɛ_S-S_ = 0.00 kT, to yield an A/B compartment strength similar to that observed in the experimentally generated PD-D2 Hi-C maps. The experimentally measured value of S-S_inter_ is shown as a purple line. (**G**) S compartment smoothness plotted as a function of compartment strength. Simulations varied in ɛ_A-A_, ɛ_B-B_, λ, d_S_, and d_A&B_. Smoothness was computed as the standard deviation of ^observed^/_expected_ S-A, S-B, and S-S_inter_ contacts. The experimentally measured value of S smoothness from the PD-D2 maps is shown as a purple line. (**H**) Comparison of the experimental data for chromosome 15: 0-6,500,000 (mapped with distiller) versus the best polymer model (ɛ_A-A_ = 0.16 kT, ɛ_B-B_ = 0.08 kT, ɛ_S-S_ = 0.00 kT, λ = 55 kb, d_S_ = 19 kb, d_A&B_ = 190 kb). *Left:* Experimentally generated Hi-C (*top, right half of the map*) versus the *in silico* generated Hi-C map from the best model (*bottom, left half of the map*). *Right*: Summary statistics (^observed^/_expected_ off-diagonal compartment and on-diagonal S_intra_ enrichments) for the experimentally generated data (*left*) and best model (*right*). (**I**) Parameter values for extruder properties (*left*) and homotypic A-to-A and B-to-B affinities (*right*) of the successful models, sorted by rank of their mean square error of their summary statistics compared to those of the experimental data. All models lacked S-to-S affinities.

We first reaffirmed our key findings from the minimal model that neither phase separation alone (Figure 4B) nor with uniform extrusion (TAD-like) (Figure 4C) can generate secluded S domains. For phase separation-only models, we varied A and B homotypic affinities in the absence of loop extrusion, but this did not produce the desired combination of S_intra_ (feature *i*) and S-S_inter_ values (feature *ii*) (Figure 4C). With uniform extrusion, we similarly could not reproduce features *i* and *ii* quantitatively, regardless of processivity (λ) or extruder density (d_S_) (Figure 4C). For the targeted loading models, some, but not all, models quantitatively reproduced experimental values for features *i* and *ii*. We noticed that different aspects of extrusion dynamics influenced either feature *i* or feature *ii* (Figure 4D). Specifically, the *relative* abundance of extruders in S versus A and B (d_A&B_/d_S_) appeared to have a strong effect on feature *i*. Likewise, the overall abundance of extruders in S (separation in S, d_S_) influenced feature *ii* (Figure 4D, arrows). These observations prompted us to more deeply investigate how different parameters of our model control the different features of S.

We first asked how such strong domains of S_intra_ contacts (feature *i*) arise from targeted loading. We systematically varied processivity (λ), mean extruder separation in S (d_S_), and mean separation in A and B (d_A&B_) independently. Consistent with Figure 4D, our simulations revealed that feature *i* is driven by the *relative* abundance of loop extruders in S (d_A&B_/d_S_) and is insensitive to processivity (Figure 4E). For a given set of compartment affinities (A-to-A affinity = 0.12 kT, B-to-B affinity = 0.04 kT), achieving the desired level of relative compaction (S_intra_ = 1.78, feature *i*) required extruders to be approximately ten-fold more abundant in S compared to A and B (i.e., d_A&B_/d_S_ ≈ 10-15, Figure 4E). Increasing A/B compartment strength allowed relative abundance in S to decrease slightly to achieve the desired S_intra_ enrichment (d_A&B_/d_S_ ≈ 5-10, Figure S6A). This suggests that S compaction results from substantial recruitment of extruders to S, comparable to targeted cohesin loading in other contexts ^43^.

We next uncovered that S-to-S compartment attenuation (feature *ii*) is predominantly controlled by the overall abundance of extruders in S (d_S_) (Figure 4F). This contrasts with vertebrates, where extrusion modulates compartment strength by both extruder processivity (λ) and separation (d). Because S domains are small (median 75 kb for our simulated region), even modest extruder processivities (≥ 56 kb) generate high loop coverage in S (Figure S6B). Therefore, extruder abundance governs S compaction (Figure S6C) and in turn S-S_inter_ seclusion (Figure 4F). To generate distal S-S_inter_ depletions similar to those measured in the Hi-C, the abundance of extruders in S must be high (small d_S_). Based on the modeled region, where the experimentally measured ^observed^/_expected_ S-S_inter_ depletion (feature *ii*) is 0.7, we estimate extruder separation (d_S_) is 20-40 kb (Figure 4F). A precise estimate depends on the strength of A/B compartmentalization, as stronger compartmentalization can promote S-S_inter_ contacts (i.e., loss of feature *ii*, Figure S6D).

We next sought to explain the most notable feature in *B. mori* Hi-C maps: S smoothness (feature *iii*). For smooth lanes of contact depletion, S must be similarly depleted in contacts with A, B, and other S domains. While S-S_inter_ depletion (Figure 4F) is primarily controlled by extrusion in S (d_S_), loop extrusion in S has only a minor effect on S-A and S-B contacts (Figure S6E). Rather, S-A and S-B contacts are predominantly controlled by A and B phase separation: loss of S-A and S-B contacts reflect the demixing (phase separation) of A and B, respectively (Figure S6E). Therefore, to achieve similar levels of S-A and S-B contacts, the level of phase separation of A and B must be approximately equal (i.e. ^observed^/_expected_ A-A contacts ≈ ^observed^/_expected_ B-B contacts). Additionally, when A/B compartmentalization strengthens, smoothness is also lost because S-S_inter_ increases. Therefore, A/B compartmentalization must be low to maintain smoothness (Figure 4G, Figure S6E and S6F).

We find *B. mori* chromatin organization arises from a unique interplay of multiple parameters, allowing us to estimate model parameters (d_S_, d_A&B_/d_S_, A-to-A, and B-to-B affinities) from the different Hi-C features of S (Figures 4E-G). To develop a quantitatively accurate *B. mori* chromosomal model, we integrated our estimates and insights from Figures 4E-G, sweeping over estimated extrusion and compartment parameters. Models with high levels of targeted extrusion (such as d_S_ = 19 kb, d_A&B_ = 190 kb; d_A&B_/d_S_ = 10) and equally modest A and B compartmentalization reproduced Hi-C maps qualitatively and ^observed^/_expected_ enrichments quantitatively (Figures 4H, 4I). Other successful models also had these characteristics (Figure 4I). As predicted, models with any parameter outside of estimated ranges (increasing d_S_, reducing d_A&B_, high or unequal A-A and B-B contact strength) systematically caused models to fail.

Our models reveal a previously uncharacterized interplay of loop extrusion and chromatin compartmentalization in shaping the genome. Loop extrusion in *B. mori* is targeted to S domains, which leads to higher S compaction and suppression of (off-diagonal) compartment contacts between S. Together, these mechanisms generate the striking chromatin contact patterns that dictate 3D genome architecture. We next used our *B. mori* chromosomal model to make predictions on spatial localization.

### S domains are preferentially found on the surfaces of chromosome territories

The spatial partitioning of active and inactive chromatin within the nucleus has previously been attributed to conventional compartmentalization ^33^. We asked whether the seclusion of compartment S confers distinct spatial positioning using our best-performing model of chromosome 15 (Figure 4H). By analyzing the radial positioning of A, B, and S, we found a strong preference for S domains to localize to the periphery of chromosome territories (CTs) (Figures 5A, 5B). A and B both showed preference for localizing to the center of the territory, with A being more centrally biased. When compared to an analogous model without extrusion, S was more interspersed throughout the territory. Models with higher targeting of extrusion to S (no loading in A and B) showed even stronger radial preference of S. Therefore, our models predict that extrusion in S modulates its spatial localization, biasing it to the periphery of CTs.

**Figure 5:**
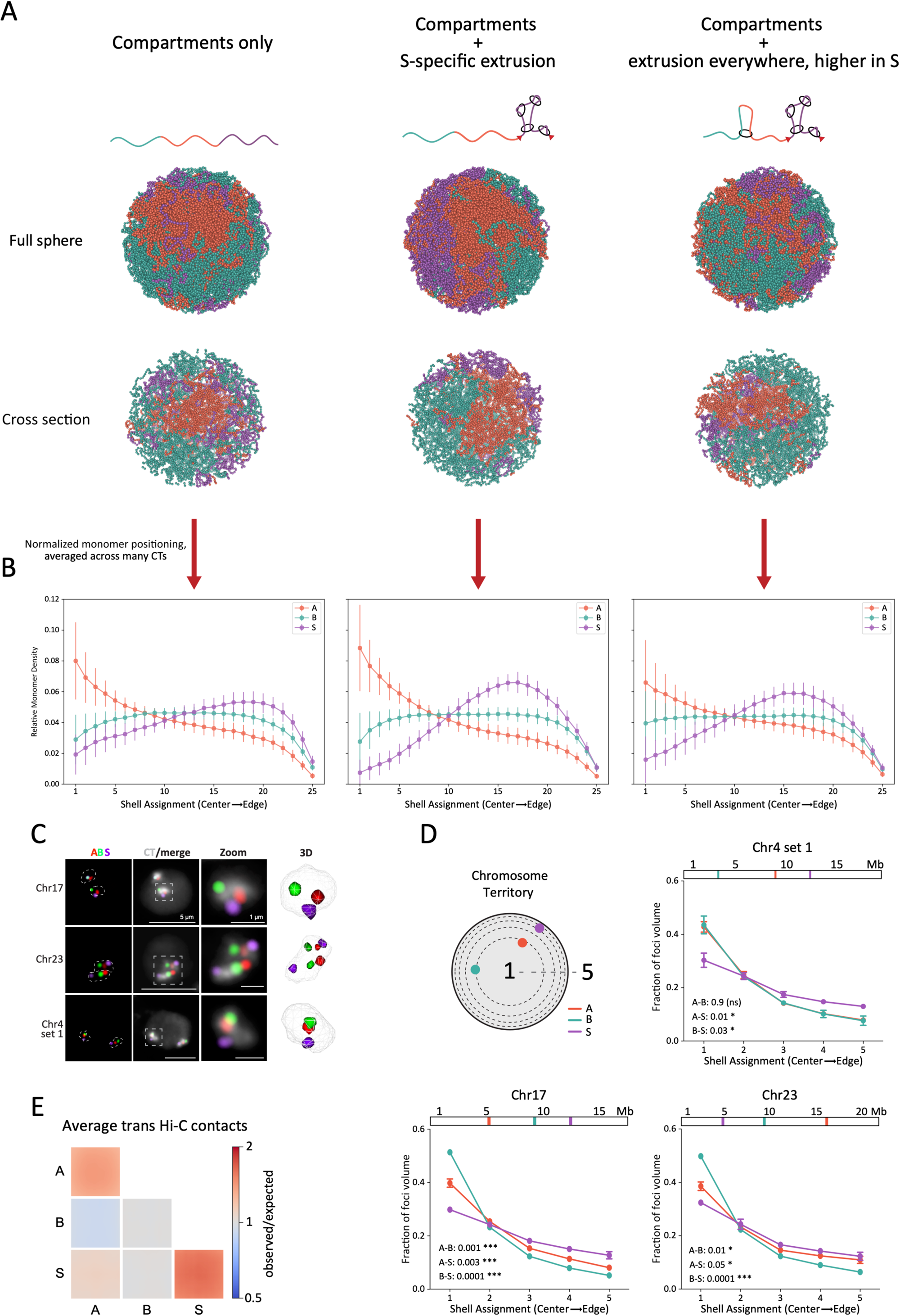
Compartment S is peripherally located within chromosome territories, with modeling indicating a role for loop extrusion in this positioning. (**A**) Example conformations for three models of chromosome 15, with the top row’s renderings reflecting the entire spherically confined simulation and the bottom row’s renderings representing cross sections through the center of the sphere. Each column represents a model introducing a new source of organization to the chromosome; (*left*) a rendering of a phase-separation-only model (ɛ_A-to-A_ = 0.16 kT, ɛ_B-to-B_ = 0.08 kT, ɛ_S-to-S_ = 0.00 kT); (*middle*) a rendering of the same compartment model with loop extrusion within S only (λ = 55 kb, d_S_ = 19 kb); (*right*) a rendering of the same compartment model as the far left but with different loop extruder separations in S versus A and B (λ = 55 kb, d_S_ = 19 kb, d_AB_ = 190 kb; the best model from the previous section). (**B**) Relative monomer densities for A, B, and S monomers for the three models detailed in Panel 5A. (**C**) Four-color Oligopaint FISH labeling single A (red), B (green), and S (purple) domains as well as the corresponding CT (white). *First column*: Oligopaint labeling of domains, with white dashed lines indicating CT edges. *Second column*: Oligopaint labeling of domains merged with Oligopaint labeling of CTs. *Third column*: zoom-in views corresponding to boxes traced in column two. *Fourth column*: 3D rendering of zoomed CT from TANGO ^91^. Microscope images are Z projections of 10 Z stacks. The background in the CT channel acts as a proxy for the nuclear edge. (**D**) Shell analysis measuring compartment foci positions within their CTs for Chr04, 17, and 23. The location of Oligopaint FISH probes within the chromosome is shown above each plot. Dots indicate means of 3 biological replicates (different embryos, n > 250 nuclei). Error bars show standard error of the mean. P-values were generated from unpaired t-test (Kolmogorov-Smirnov test) between distributions and are indicated at the bottom left of each graph. (**E**) Average ^observed^/_expected_ *trans* (inter-chromosomal) contacts within and between all A, B, and S compartments.

To test these findings, we used Oligopaint FISH to label portions of single A, B, or S domains as well as the whole CTs for Chr04, Chr17, and Chr23 in embryonic nuclei (Figures 5C, S7B and S7C). Shell analysis (Material and Methods) revealed that S domains are more likely to occupy peripheral CT shells compared to A and B domains (Figure 5D). In addition, measuring the distance from the domain center to the CT edge showed that S domains are closer to the CT edge than A or B domains for all chromosomes and loci analyzed (Figure S7D). These observations were confirmed by additional experiments with super-resolution microscopy (Figure S7E and S7F).

Although inter-chromosomal contacts were sparse in the Hi-C data, we asked whether this peripheral localization could influence the average contact frequency among S domains in *trans*. Therefore, we computed the average *trans* ^observed^/_expected_ contacts between each compartment type (Figure 5E). We found that the average value of S-S_inter_ *trans* contacts is higher compared to any other combination in *trans*, and, in particular, A-A and B-B contacts. This is consistent with the preferential positioning of S towards the periphery of CTs, a favored location for *trans* contacts.

Our DNA FISH reveal S domains are preferentially located at the CT peripheries, which is further supported by the Hi-C. Our modeling indicates that this may be caused by a previously unknown effect of loop extrusion: its ability to influence the spatial positioning of chromatin in the context of its chromosome territory.

### Compartment S changes during development

Taking advantage of our Hi-C datasets from different developmental stages, including three embryonic stages and one adult stage, we next explored the developmental dynamics of compartment S. Based on initial visual inspections, we could identify domains switching to or from S on multiple chromosomes between timepoints (see examples in Figure S8A). To test the dynamics of S compartment switching more systematically, we repeated the compartment calling protocol for the adult stage (Adult Heads, AH). We found that a comparable fraction of the AH genome (12%) folds into compartment S, compared to the PD-D2 embryonic stage (Figure S8B). Nevertheless, several embryonic S domains visually show checkerboard patterning in the AH Hi-C maps, including three of the largest S domains on Chr06 and Chr23 (Figures 6A and S8C). To compare the two datasets further, we restricted our analyses to large (>200 kb) domains to allow for visual confirmation of compartment assignment based on the Hi-C contact pattern. We found that about 45% of S domains defined in the PD-D2 embryonic stage are maintained in the AH (referred to as S->S) (Figure 6B). This fraction is much lower compared to that of the A or B domains. Large S domains that change compartment assignment in the AH exclusively turn into B domains (referred to as S->B). Notably, this change coincides with a significant reduction in gene expression levels (Figures 6C and S8D). The Hi-C contact maps at S domains that turn into B show a loss of features associated with loop extrusion, including insulation points and off-diagonal dots (Figures 6A and S8C). Weakening of insulation is also evident when comparing pileup contact enrichments of boundaries called within large S domains that are maintained (S->S) or lost (S->B) in the AH data (Figure 6D). The loss of extrusion features and intra-domain compaction, together with the increased checkerboard patterning of S domains that turn into B, are consistent with our model that targeted loop extrusion underlies the spatial segregation of compartment S by counteracting compartmentalization.

**Figure 6:**
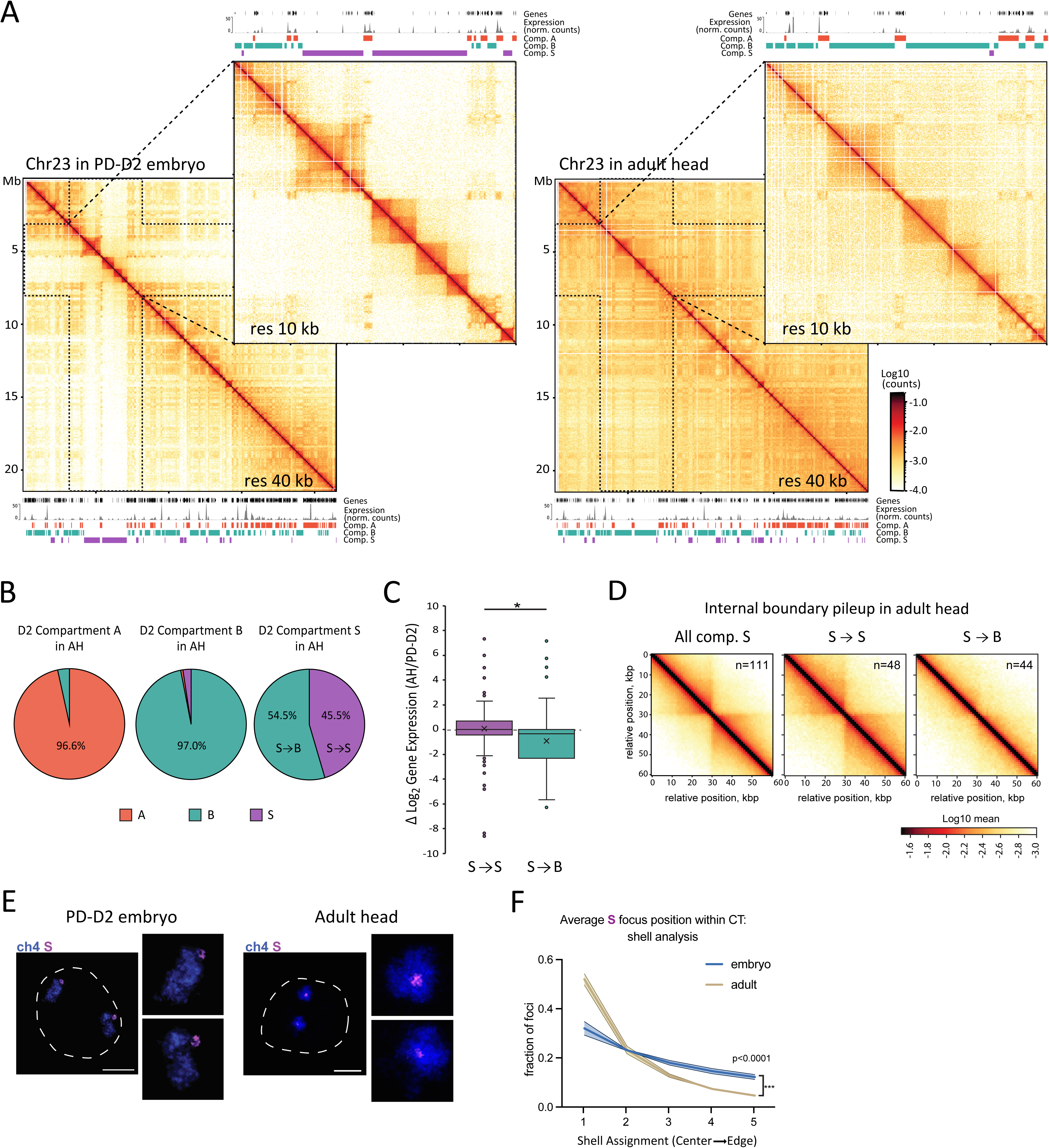
Compartment S changes during development. (**A**) Hi-C contact maps of the full Chr23 at 40 kb resolution and Chr23:3,000,000-80,000,00 region at 10 kb resolution in PD-D2 embryos on the left and in adult head (AH) on the right. Below or above each matrix are gene locations, RNA-seq expression aggregated at 1 kb resolution and compartments A, B, and S assignments, for the corresponding stage and region. (**B**) Compartment assignment in AH datasets of large domains (>200 kb) assigned to large compartment A, B, or S in PD-D2, in % of the compartments assigned in PD-D2. (**C**) Box plots of distributions of log_2_ ratio of gene expression (TPM) between PD-D2 embryos and AH. In purple is the distribution for genes that are in domains assigned to compartment S in the two stages (S◊S) (n=119), and in green is the distribution for genes that are in region assigned to compartment S in PD-D2 and that switched to B in AH (S◊B) (n = 120). Boxed region corresponds to data between first to third quartile. Line indicates median of distribution while cross corresponds to mean. Whiskers extend to lowest and highest non-outlier data points, and dots outside correspond to outliers. Asterisks indicate distributions significantly different by Kolmogorov-Smirnov test (*: P-value = 0.01 - 0.05). (**D**) Pileup plots centered on internal insulation points within all large S compartment domains (>200 kb, excluding 40 kb on each side of domain boundaries) in AH or S◊S and S◊B categories, as described in the right panel of (B), at 5 kb resolution and extending to 30 kb on each side. (**E**) Two-color STED labeling chr04 CT and single S domain in cells from PD-D2 embryos (left) or adult heads (right). Chr04 is shown in blue, and the S compartment domain is shown purple. Dashed line indicates nuclear edge as determine by confocal microscopy. (**F**) Shell analysis measuring S domain positions within Chr04 in embryos (blue) and adult heads (tan). Mid line = average of 2-3 biological replicates (n>30 nuclei each). Error bars (light shading) show standard error of the mean. P-values were generated from unpaired t-tests (Kolmogorov-Smirnov tests) between distributions and are indicated at the bottom right of the graph.

To test whether the change in compartment identity also translate to a change in domain position within CTs, we took advantage of the fact that one of the three S domains that we chose for Oligopaint FISH analyses turned into B in the AH Hi-C dataset. Consistent with the change in compartment identity, super-resolution microscopy revealed that the same region on Chr04 localizes more internally in its CT in nuclei of the adult *B. mori* head compared to embryos (Figure 6E and F). In contrast, a B domain used as a control remained unchanged between the two stages (Figure S8E and F). These results were further validated using wide-field microscopy, labeling A, B, and S domains as well as the entire CT on Chr04 in AH. Notably, in AH cells, the S domain that transitioned into B no longer exhibited a tendency to be positioned closer to the CT periphery (Figure S8G and H). The developmental dynamics of several S domains argue against a strict genetic specification of compartment S but rather support the presence of epigenetic features involved in its formation.

## Discussion

Our investigation of *B. mori*’s genome organization reveals both conserved principles and novel genome-folding behaviors. As seen in other eukaryotes, we observe the formation of CTs and spatial segregation of chromatin into active A and inactive B compartments. However, unlike other eukaryotes, we have observed much stronger chromosome territoriality and a novel type of compartment, S, which lacks the characteristic checkerboarding of A/B compartments and appears to be rich in loop extrusion activity. Our modeling indicates that a unique interplay of chromatin compartmentalization and loop extrusion may drive the formation of these spatially secluded domains that appear to change throughout development.

The remarkably strong CTs and the low frequency of inter-chromosomal contacts are consistent with a recent whole-chromosome Oligopaint study of six *B. mori* chromosomes, which likewise revealed highly spatially distinct CTs ^73^. Folding, volume, and intermixing of CTs have been associated with condensin II ^64,94^. Condensin II subunits are present in the *B. mori* genome ^95,96^ and it would be interesting to evaluate whether the substantial degree of territoriality is caused by uniquely high activity of condensin II in *B. mori*. CT strength is also intriguing from an evolutionary point of view. Previous studies in *D. melanogaster* cell lines ^97^ and across human cancers ^98^ have described an inverse relationship between the frequency of inter-chromosomal contacts and the incidence of genomic translocations. Interestingly, karyotypes and synteny are highly conserved across Lepidoptera, including *B. mori*. We therefore hypothesize that the pronounced CTs may contribute to low structural variation and high karyotype conservation in these organisms by preventing structural rearrangements during mitotic divisions of cells destined to form the future germline.

The presence of compartment S, with domains that strongly self-interact but segregate from the rest of their chromosome, is remarkable. To our knowledge, there is no precedent for a compartment with contact or epigenetic profiles similar to compartment S. While other compartment types beyond A and B have been detected ^40,82,99^, they typically represent sub-types of A and B and show preferential homotypic contacts ^82^. Although the previously described “intermediate” compartment I ^27,100,101^ shares some characteristics with compartment S, such as H3K27me3 enrichment and developmental plasticity, they differ in their most prominent features. Compartment I is *enriched* in contacts with A, B, and I. A recently identified sub-compartment in HCT116 cells, termed B_0_ ^82^, likewise displays relatively low levels of compartment contrast (i.e., its compartments are smoother and they checkerboard less than other compartment types), though it does preferentially interact with other B_0_ domains. Domains of compartment S, in contrast to I or B_0_, are homogenously *depleted* in contacts with any other domain. This pattern, plus their gene composition and distinct epigenetic makeup, make S unique compared to any previously identified chromatin structure.

The formation of compartment S appears to require an interplay of mechanisms, distinct from any other described organism. While conventional compartmentalization is believed to rely on affinity between regions that share epigenetic composition ^102^, we show that affinity-mediated phase separation cannot lead to the formation of compartment S. Instead, we propose that compartment S is formed by loop extrusion activity targeted to S domains. Targeting extrusion to S domains selectively compacts them and suppresses their compartmentalization. This loop extrusion behavior is different than in vertebrates, where loop extruding cohesin (during interphase) and condensin (in metaphase) are believed to load mostly uniformly across the genome, without preference for a specific compartment type ^28,82^. Such localization of extruders to many broad epigenetic domains (tens to hundreds of kb each) has not been identified in other systems.

Targeted SMC loading to certain genetic elements is well-described across various biological systems. In the bacterium *B. subtilis*, condensins are loaded at *ParS* sites by the ParB DNA-binding protein ^103,104^. In yeast, it has been suggested that sequence context antagonizes SMCs from centromere binding ^105^. In *C. elegans*, direct loading of condensin^DC^ to specific sequences (rex sites) direct the SMC-containing dosage compensation complex to the sex chromosome ^65,106^. Most recently, targeted loading of SMCs to enhancers has been shown to form fountains or jets in Hi-C maps in *C. elegans* ^44,47^ and vertebrates ^42,43,107^. Such targeted loading in these systems may be guided by differences in DNA accessibility, sequence-specific DNA binding proteins ^103,104^, or via enhancer-specific histone marks ^42,43,107^. We speculate that such factors may target loop-extruding complexes to compartment S in *B. mori*.

In the context of targeted loading of SMC complexes, the enrichment of H4K20me1 in compartment S is interesting. In *C. elegans*, it has been shown that the SMC-based dosage compensation complex enriches H4K20me1 on the inactive sex chromosome by means of a condensin-associated demethylase ^86^. H4K20me1 is likewise deposited during the process of silencing and compacting the X chromosome in mice ^108^. Compartment S shares similarity to inactivated sex chromosomes by virtue of being locally compacted and isolated from the rest of the genome, while it is distinct from inactivated sex chromosomes by virtue of its permissiveness to gene expression. Whether enrichment of H4K20me1 aids in chromatin compaction, and what its relationship is to loop extrusion, is unclear for *B. mori*.

In addition, our analyses also support a previously unknown consequence of loop extrusion. Not only does loop extrusion activity lead to the formation of dense and secluded S domains, but it also drives their peripheral localization within CTs. This effect of loop extrusion is a novel finding, raising the possibility that loop extrusion may similarly affect large-scale spatial organization in other organisms or contexts. The underlying physics by which extrusion can achieve this architecture is yet to be understood.

The function of compartment S and the role of loop extrusion there remain intriguing questions. First, in view of the holocentric architecture of *B. mori* chromosomes, we consider the possibility that compartment S is involved in centromere specification to be unlikely. This conclusion is guided by our previous findings in *B. mori* cell lines showing that kinetochore assembly occurs in regions anticorrelated with transcriptionally active chromatin ^109^ - an epigenetic landscape that is more consistent with compartment B than S. Instead, the isolated genomic environment of S domains might ensure the precise transcriptional regulation of the genes that they contain. Our finding that S-located genes are functionally enriched in transcription-related processes might suggest that S represents a developmental transition state to either A or B, as hypothesized for compartments I and B_0_ ^82,100,101^. Such a model is supported by the observation that many S domains are variable among different developmental stages. Furthermore, S domains may represent development-control units, such as the Hox cluster, which comprises a large S domain in *B. mori* embryos (Figure S4). By analogy to the Hox cluster, where loop extrusion appears to be key to the precise sequence of gene activation ^110^, compartment S may recruit a high density of extruders to achieve precisely timed activation of genes during development. Our observation of developmental plasticity of S domains supports this hypothesis.

In summary, our study describes the unique genome organization of *B. mori.* This non-model organism has an exceptional degree of chromosome territoriality and reveals a striking new genome folding structure. Our modeling indicates that this novel structure, compartment S, can be formed by loop extrusion targeted to a specific chromatin type. Genome organization in *B. mori* thus demonstrates that the interplay of two major genome-folding processes, loop extrusion and compartmentalization, can generate unique and unexpected patterns. These findings open a new line of questions for how evolutionarily conserved mechanisms may interact differently in different organisms, as well as how changes in the epigenetic landscape during development can guide loop extrusion to reshape 3D genome architecture.

## Supporting information

GIl_et_al_SOM

## Acknowledgements

We would like to thank the members of the Lei, Mirny, and Drinnenberg labs, as well as Job Dekker and Axel Cournac for their helpful discussions. We thank Max Imakaev for extensive, critical feedback on the manuscript. We thank all members of the Koszul lab and in particular Pierrick Moreau for helping in the initial development of the Hi-C protocol. We thank Stéphane Descorps-Declère, Ilya M. Flyamer, and Nicolas Servant for their help in using bioinformatics tools. We would also like to thank Life Science Editors for editing services (www.lifescienceeditors.com). EN is funded by the National Science Foundation’s Graduate Research Fellowship. This work of LAM is supported by NIH GM114190 and NSF-ANR Program (ANR-22-CE95-0003). LAM is also a recipient of the « Chaires Internationales de Recherche Blaise Pascal » and the CNRS Fellow-Ambassador award. HM and IAD receive salary support from the CNRS. This work is supported by the Labex DEEP ANR-11-LABX-0044 part of the IDEX Idex PSL ANR-10-IDEX-0001-02 PSL, an ATIP-AVENIR Research grant, the FRM and Fondation Schlumberger (FSER202202015420), Institut Curie, the ERC (CENEVO-758757), and INSB «Diversity of biological mechanisms». This work was supported in part by the Intramural Program of the National Institute of Diabetes and Digestive and Kidney Diseases, National Institutes of Health (NIDDK; DK015602 to E.P.L.), the Eunice Kennedy Shriver National Institute of Child Health, and Human Development, National Institutes of Health (NICHD; 1K99HD104851 to L.F.R).

